# Exploring Liquid-Liquid Phase Separation in the organization of Golgi Matrix Proteins

**DOI:** 10.1101/2023.07.21.550027

**Authors:** Luis Felipe S. Mendes, Carolina G. Oliveira, Emanuel Kava, Antonio J. Costa-Filho

## Abstract

The Golgi apparatus is a critical organelle in protein sorting and lipid metabolism. Characterized by its stacked, flattened cisternal structure, the Golgi exhibits distinct polarity with its *cis*- and *trans*-faces orchestrating various protein maturation and transport processes. At the heart of its structural integrity and organization are the Golgi Matrix Proteins (GMPs), predominantly comprising Golgins and GRASPs. These proteins contribute to this organelle’s unique stacked and polarized structure and ensure the precise localization of Golgi-resident enzymes, which is crucial for accurate protein processing. Despite over a century of research since its discovery, the Golgi architecture’s intricate mechanisms still need to be fully understood. Here, we demonstrate that GMPs present a significant tendency to form biocondensates through Liquid-Liquid Phase Separation (LLPS) across different Eukaryotic lineages. Moreover, we validated experimentally that members of the GRASP family also exhibit a strong tendency for LLPS. Our findings offer a new perspective on the possible roles of protein disorder and LLPS of GMPs in the Golgi organization.

## Introduction

Eukaryotic cells evolved to possess efficient secretory machinery capable of transporting a significant fraction of their proteome (1). Proteins entering the secretory pathway are efficiently sorted to a specific destination: the extracellular space, the plasma membrane, or the endomembrane system. The machinery also has efficient quality control, avoiding the transport of misfolded proteins or complexes (2). Proteins destined for this classical secretory pathway are synthesised in the endoplasmic reticulum (ER) interior, where they are first folded and possibly modified by the post-translation ER machinery (2). Subsequently, these secretory cargoes are transferred to the ER exit sites (ERES), where they are assembled into secretory vesicles coated with coatomers (COPII), routed for the ER-Golgi intermediate compartment (ERGIC), and later to the *cis*-Golgi face (3, 4).

The Golgi apparatus, a central organelle in eukaryotic cells, plays a pivotal role in trafficking, processing, and sorting proteins and lipids. This membrane-bound organelle exhibits a unique architecture consisting of a stack of flattened cisternae interconnected by tubular-vesicular networks. The cisternae can be further classified into three distinct regions: *cis*, *medial*, and *trans*, each responsible for specific functions in protein processing and glycosylation. The maintenance of this intricate structure is crucial for the Golgi’s function and relies on a diverse group of proteins, collectively known as the Golgi matrix proteins (GMPs). In the organisation of the Golgi stacks and cisternae formation, it was observed that in cells treated with brefeldin A (an inhibitor of COPI-mediated transport from the Golgi to the ER), the Golgi membrane and its integral proteins relocated to the ER (5). On the other hand, the GMPs accumulated in the cytoplasm formed a ribbon-like reticulum in the correct Golgi location (5). Based on this observation, it seems that GMPs could generate the higher-order architecture of the Golgi by a mechanism independent of other Golgi proteins. Although an orchestrated action involving all the GMPs provides the total adhesive energy dictating Golgi cisternal stacking (6), how these proteins act in the general organisation/dynamics remains unknown.

The Golgins and the Golgi Reassembly and Stacking Proteins (GRASP) are the main protein families composing the GMPs. The Golgins are predominantly extended coiled-coil proteins predicted to reach an extension between 100-600 nm (7, 8). They attach to the Golgi cisternae through either their carboxy terminus, directly interacting with the membrane surface (7) or a membrane-associated partner (9). Golgins participate in vesicular traffic, Golgi maintenance and dynamics, and the positioning of the Golgi apparatus within cells (8, 10). On the other hand, GRASPs constitute a family of peripheral membrane proteins essential for general Golgi organisation into the ribbon-like structure (9) and Golgi integrity for the correct protein sorting and glycosylation along the secretory pathway (11). GRASPs are observed in all branches of the eukaryotic tree of life, although not equally spread in all supergroups (12). GRASPs are collapsed intrinsically disordered proteins (IDPs) with high promiscuity for mediating protein-protein interactions with several different protein partners (13–15). Their unusual structural plasticity (16) might be the converging feature of GRASPs in multiple cell functional tasks, including their direct participation in Golgi dynamics during mitotic and apoptotic times (17–19) and their self-association during stress conditions triggering unconventional protein secretion pathways (20–23). GRASPs interact with the membrane via a lipidation of its Gly2 using a myristoyl-switch mechanism (24, 25). They also anchor some Golgin representatives, including GM130, Golgin45, and p115 in mammalian cells and the Golgin BUG1 in Yeast (9, 26). This anchoring is essential for the correct Golgi organisation (9, 27).

In 2019, James Rothman suggested that the Golgi, akin to the nucleolus, would fundamentally be liquid with a phase-separated internal organisation (28). This suggestion came after discovering that many human Golgins can efficiently populate a liquid-like droplet state once they are overexpressed *in vivo* and dispersed in the cytosol (29). In recent years, liquid-liquid phase separation (LLPS) was discovered to form and organize biomolecular condensates within cells. LLPS is a process by which a homogenous solution of macromolecules separates into distinct liquid phases, often leading to the assembly of membrane-less compartments with unique biophysical properties (30). This phenomenon has been implicated in cellular functions, such as signal transduction, RNA processing, and stress response (31–33).

Here, we build on the discussion of the role of LLPS in regulating cellular processes by expanding its potential to describe several characteristics of the Golgi apparatus, especially its dynamic nature and unusual flattened membrane organisation. It is important to note that the Golgi is a membrane-bound organelle, setting it apart from membrane-less organelles like the nucleolus. Furthermore, any LLPS at the Golgi involving Golgi-matrix proteins would need to occur on a membrane surface. The remodelling of cell membranes requires their bending. Membrane bending is an essential cellular process with implications in numerous aspects of cell biology, such as membrane traffic, cell motility, organelle creation, and cell division (34–36). Besides the traditional bending induced by well-shaped proteins, it has been recently shown that protein phase separation occurring on the surfaces of both synthetic and cell-derived membrane vesicles induces significant compressive stress within the membrane plane, which prompts the membrane to bend inward, forming protein-lined membrane tubules (35, 37). In this case, the proteins undergoing phase separation contain substantial regions of intrinsic disorder, which do not possess a stable three-dimensional structure (37, 38). Fascinatingly, disordered regions have recently been observed to form networks sustained by weak, multivalent contacts, facilitating the assembly of protein liquid phases on membrane surfaces (39, 40).

Since LLPS can affect membrane bending dramatically (35, 39, 41) and some Golgins have been shown to undergo LLPS in cells (29), it would be reasonable to argue that there might be a direct correlation between the LLPS-propensity of GMPs and the unusual Golgi cisternae shape. However, it is unclear how the LLPS propensity spread among the GMPs and how this is conserved in the remaining Eukaria tree. Here, we highlight the significant prevalence of intrinsically disordered regions within GMPs across Eukarya and how these regions are linked to a high propensity for those proteins to form liquid-like condensates via LLPS. We also describe a detailed experimental study of LLPS of human GRASP55 and GRASP65 to confirm some of our predictions.

## Results and Discussions

Despite the commonly accepted presence of coiled-coil regions within Golgins, this still lacks experimental validation. There has also been a limited investigation into the prevalence of structural disorder in these proteins. Coiled-coil regions are frequently predicted as disordered sequences at rates similar to genuine intrinsically disordered segments identified in DisProt (42, 43). Although coiled coils often transition from a disordered to an ordered state upon assembly, they hardly ever evolve into highly compact structures. Their extended configuration and the presence of extensive multivalent surfaces possibly underlie the observed propensity of specific coiled-coil-containing proteins to undergo LLPS in cells (29, 44, 45). This behaviour resembles the multivalency characteristic of disordered proteins, enabling many of them to engage in numerous weak interactions concurrently. Such a feature significantly enhances their likelihood of undergoing LLPS. Despite these insights, a thorough large-scale investigation remains lacking, particularly one that evaluates GMP disorder and quantifies amino acid residues conducive to more compact structures.

We started by exploring the structural properties of the metazoan GMPs. Identifying all GMPs in metazoans is an active research area, and the comprehensive list has not been definitively established. An extra difficulty is that different organisms can express different sets of Golgins. We selected some well-known GMPs and associated proteins that are known to play a role in maintaining Golgi structure and function. It is important to emphasise that we might not have exhausted the full list of GMPs. Some GMPs commonly identified in metazoans are: GM130 (GOLGA2), Golgin-45 (BLZF1), Golgin-160 (GOLGA3), AKAP9 (CG-NAP), Giantin (GOLGB1), GCP60 (ACBD3), GCC88 (GCC1), Golgin-97, Golgin-245 (GOLGA4), SCYL1BP1 (GORAB), GCC185 (GCC2), Bicaudal-D (BCD1 and BCD2), NEEC1 (JAKMIP2), GRASP65 (GORASP1) and GRASP55 (GORASP2). These families form the list to be discussed now in the text. Other important GMPs and Golgi structure-associated proteins like CASP (GOLGA8), Golgin-84 (GOLGA5), p115 (also known as USO1), Zinc Finger Protein-Like 1 (ZFPL1), and GMAP-210 (also known as TRIP11) are also found in metazoans but have a broader presence in the Eukary tree, and so will be discussed later in the text in a broader eukaryotic perspective. It is important to note that different proteins can have multiple names or synonyms, adding to the complexity of cataloguing them all.

Not all the proteins listed in the previous paragraph are Golgins. Golgins are typically predicted to have coiled-coil regions; many have a C-terminal GRIP domain or a similar structure. In the previous list, proteins like GRASP65 and GRASP55, while important to the function and structure of the Golgi, are not classified as Golgins. The same is true for the ZFPL1. The distinction between Golgins and other Golgi-associated proteins is not always clear, and there may be exceptions or debates over how specific proteins are classified. The pervasive presence of Golgins with their long rod-like structures establishes an extensive ribosome-free zone around the Golgi apparatus, a feature readily observable in microscopy images (46).

Our comprehensive analysis spanned >28,000 different protein sequences and employed metrics to shed light on the proteins’ inherent tendency to adopt extended configurations (out of classical collapsed ones), consequently exposing their multivalent properties. Such multivalence is pivotal for LLPS and a central theme of our investigation. We started utilising the Hydrophobicity versus Net Charge analysis to address this issue. Initially proposed by Uversky, this method provides a binary classification of proteins (47). The method builds a clear boundary (red line in Figure S1) separating “natively disordered proteins” from “ordered proteins”. Predominantly used to differentiate extended IDPs from globular proteins, this analytical approach unveiled a critical trait: most GMPs exhibited behaviours akin to extended IDPs (Figure 1). This finding underscores the conservation of sequence features within GMPs, including low net hydrophobicity and high net charge.

**Figure 1:**
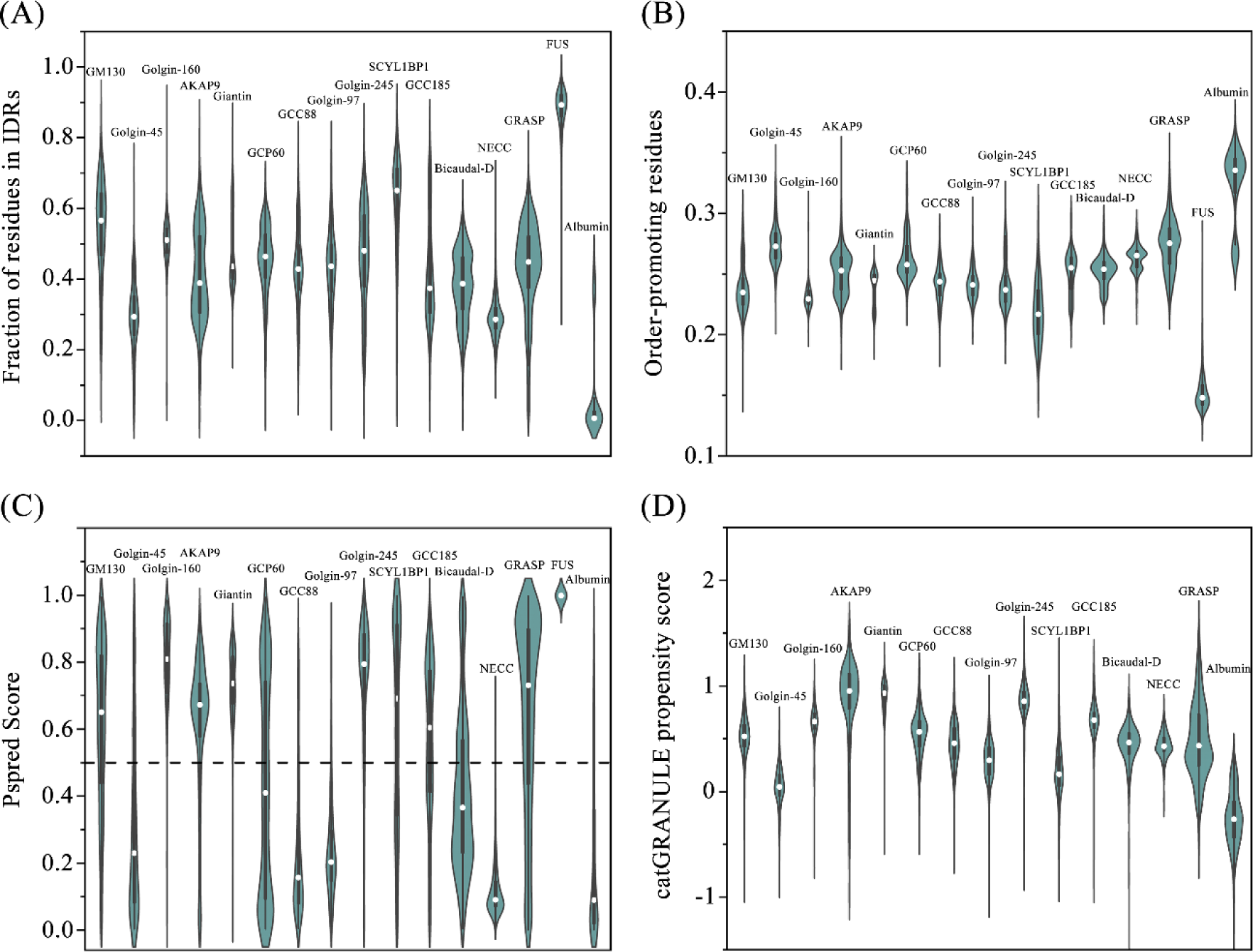
Analyses of Amino Acid Composition and LLPS-Propensity Across Metazoan GMPs. (A) Representation of residue fractions exhibiting a score exceeding the 0.5 thresholds as per IUPred2 long disorder predictions. (B) Enumeration of order-promoting residue fractions (’I’, ‘C’, ‘L’, ‘V’, ‘W’, ‘Y’, ‘F’) as defined by reference (50). (C) Application of the machine-learning-based PSPredictor (51) for LLPS prediction. A protein sequence is categorised as phase separating if the PsPred Score exceeds 0.5. (D) Assessment of LLPS propensity utilising the catGRANULE algorithm, where propensity scores are evaluated in the context of proteome distribution, as conveyed through a cumulative distribution function. A higher score corresponds to a higher propensity for a protein to phase separate. As dosage-sensitive proteins are known to score approximately 0.5 (as per reference (52)), we designate this score as our threshold in the analyses.

Notable exceptions to this trend are GRASPs and Golgin-45, which appear to reside more frequently on the “well-folded” side of the Uversky plot (Figure S1). GRASPs consist of two primary regions: the structured GRASP domain and a fully disordered C-terminus, the latter typically being the larger portion of those proteins (15). Previous molecular biophysics analyses indicated GRASPs as molten-globule proteins, suggesting an intermediate level of structural order (16, 21). Golgin-45 is the primary binding partner of mammalian GRASP55 (13, 48). Unlike GRASP, which is predicted to be present in the Last Eukaryotic Common Ancestor (LECA), Golgin-45 appears to be holozoan-specific (49). Prior research demonstrated that the progenitor of the paralogous proteins GRASP55 and GRASP65 could bind Golgin-45 with high affinity (13). In this context, GRASPs and Golgin-45 neither exhibit the mean net charge/hydrophobicity ratio characteristic of extended IDPs, like other GMPs, nor fit in the classic well-folded realm of protein structure.

To expand the previous observations, we explored the relative amount of IDRs in the protein sequences to get more quantitative details of the disordered behaviour. Drawing from the rich tapestry of protein families, we singled out two distinct families for comparison with our database. On one end of the spectrum, we considered the mammalian Albumins— representative of structurally well-ordered proteins—and on the opposite end, we included the RNA binding protein FUS, a classic representative of the extended IDPs found in vertebrates. The median of disorder fractions varied substantially from around 30% of the total protein sequence for Golgin45 and NEEC to nearly 70% for SCYL1BP1 (Figure 1A). Our data showed that most families were composed of a median ranging from 40-60%, explaining why these proteins behave more similarly to extended IDPs in the Uversky plot analyses.

To deepen our understanding of the structural propensities of the GMPs, we analysed their overall composition of order/disorder-promoting residues. In the realm of protein structure, the concepts of “disorder-promoting residues” and “order-promoting residues” refer to the propensities of specific amino acids to promote either a structured (ordered) or unstructured (disordered) conformation of the protein chain. The results observed for the GMPs tested were between the extreme situations observed for order-promoting residues in FUS (∼15%) and Albumin (∼33%), (Figure S1), again suggesting a high degree of structural flexibility. GMPs harboured fewer regions of order and exhibited higher flexibility (Figure 1B), suggesting that their function is likely linked to these structural characteristics. For example, GMPs tend to adopt extended structures, which give rise to the extensive ribosome-exclusion area surrounding the Golgi in microscopy images (46, 53).

IDPs and IDRs are unique segments of proteins that lack a fixed or stable three-dimensional structure under physiological conditions *in vitro*. They are associated with many cellular functions, including molecular recognition, molecular chaperones, protein scaffolding, flexible linkers, among others (54, 55). The flexibility and diverse binding potential of IDPs/IDRs are often associated with an increased propensity for liquid-liquid phase separation (LLPS) (40, 51). However, this does not necessarily imply a direct correlation. LLPS is like oil droplets forming in water: specific proteins or protein complexes separate from the surrounding solution to form concentrated, liquid-like droplets within the cell, and this phenomenon is now recognised to be widely spread *in vivo* (56). Predicting the propensity of a protein or a protein region to undergo LLPS is a vibrant area of investigation in current bioinformatics and biophysical research. Several computational tools are available to predict LLPS propensity, with many focusing on the presence of IDRs and sequence-based features like amino acid composition, charge, and hydrophobicity (51, 52). These include tools like PSPred and catGRANULE (51, 52).

PSPred is a machine-learning, sequence-based tool that queries and predicts potential phase separation proteins. Its output is represented by the “PSP Score”, which ranges from 0 to 1, with 1 indicating the highest propensity for LLPS (51). In contrast, the catGRANULE algorithm delivers a protein-level score as a cumulative distribution (52). It is noteworthy that catGRANULE does not prescribe a default cutoff, but based on how the model is trained, any score greater than 0 already suggests that a protein is predisposed to phase separation (52). Nevertheless, we have opted for a cutoff value of 0.5, which corresponds to the median in the distribution of scores determined in the positive training set. We utilised these two analytical tools to assess our GMP database, focusing on evaluating the propensity of LLPS and its conservation within specific protein families. We only excluded the FUS family data from the catGRANULE output analysis due to its distinct inclination towards a phase-separated state. Its median value exceeds 4, skewing the graph and complicating the comparison with other protein families (Figure S2).

Previously, metazoan-specific Golgins, such as GM130, Golgin-97, Golgin-245, GCC88, and GCC185, were observed to undergo phase separation when overexpressed *in cell* (29, 45). Interestingly, Golgin-97 and GCC88 families did not demonstrate a strong propensity towards LLPS (Figures 1C and D). Similarly, Bicaudal-D, NEEC, and Golgin45 lacked such strong propensity (Figures 1C and D). To delve deeper into these protein families with lower propensities, we tested human protein representatives using the PhaSePred Algorithm. This tool is a meta-predictor that aggregates the prediction scores of several LLPS-associated predictors (57). Notably, the human Golgin-97 protein (Uniprot Q92805) returned a Feature PS-Part score (a metric related to proteins whose phase separation behaviours are regulated by protein or nucleic acid partner components) exceeding 75%. This score points to a strong, albeit partner-dependent, propensity for LLPS. Similar patterns were detected for GCC88 (Uniprot Q96CN9), Bicaudal-D (Uniprot Q96G01), and Golgin-45 (Uniprot Q9H2G9), which produced PS-Part scores of approximately 64%, 77%, and 52% respectively. In contrast, NEEC (Uniprot Q96AA8) generated a markedly lower score, at around 32%.

Therefore, all GM protein families tested (except for NEEC) are suggested to phase-separate spontaneously or in a partner/context-dependent process. This prevalence was particularly notable, considering the unique evolutionary paths of these proteins across Eukaria and their low primary sequence conservation. The significant conservation of this physicochemical property stood out, underscoring its potentially crucial role in those proteins’ functions and interactions. Since LLPS appears inherent to metazoan GMPs, it is tempting to speculate that this property is important to the proteins’ functionality. As previously discussed, one of the principal roles of GMPs lies in organising and preserving the Golgi apparatus. However, characteristics such as flattened cisternae and stacked Golgi are not unique to metazoans.

Furthermore, considering the evolutionary pathways and conservation of certain GMPs, there is a compelling argument that Golgi stacking may have originated in the common eukaryotic ancestor. In a study grounded on phylogenetics, Barlow and colleagues identified several proteins implicated in the Golgi structure, which were inferred to be present in the Last Eukaryotic Common Ancestor (LECA). These protein families, which share roles in delineating separate Golgi structures, include GMAP, GRASP, TMF, Golgin-84, CASP, p115, GRIP-containing protein, and zinc finger protein-like 1 (ZFPL1) (49).

Therefore, we expanded our earlier analyses to encompass a database of these “ancient” GMPs, where we collected a sum of different modern orthologs from all possible branches of the Eukarya tree. This extension aimed to investigate whether the high LLPS tendency observed in metazoans is also prevalent across Eukarya. Our analyses confirm a high prevalence of protein regions with a high propensity to be intrinsically disordered, with medians ranging from ∼20% (p115) to ∼60% (TMF) of the entire protein sequence (Figure 2A). Interestingly, even other properties like pI are highly conserved across the eukarya GMPs predicted to be present in LECA (Figure 2B). In alignment with our previous findings, most GMPs also exhibited a high propensity for spontaneous LLPS, including the families of GRASP, GMAP-21, TMF, Golgin-84, and GRIP-CP (Figures 2C and 2D). The exceptions were ZFPL1, p115, and CASP, with scores as low as those observed in our control dataset of Albumins.

**Figure 2:**
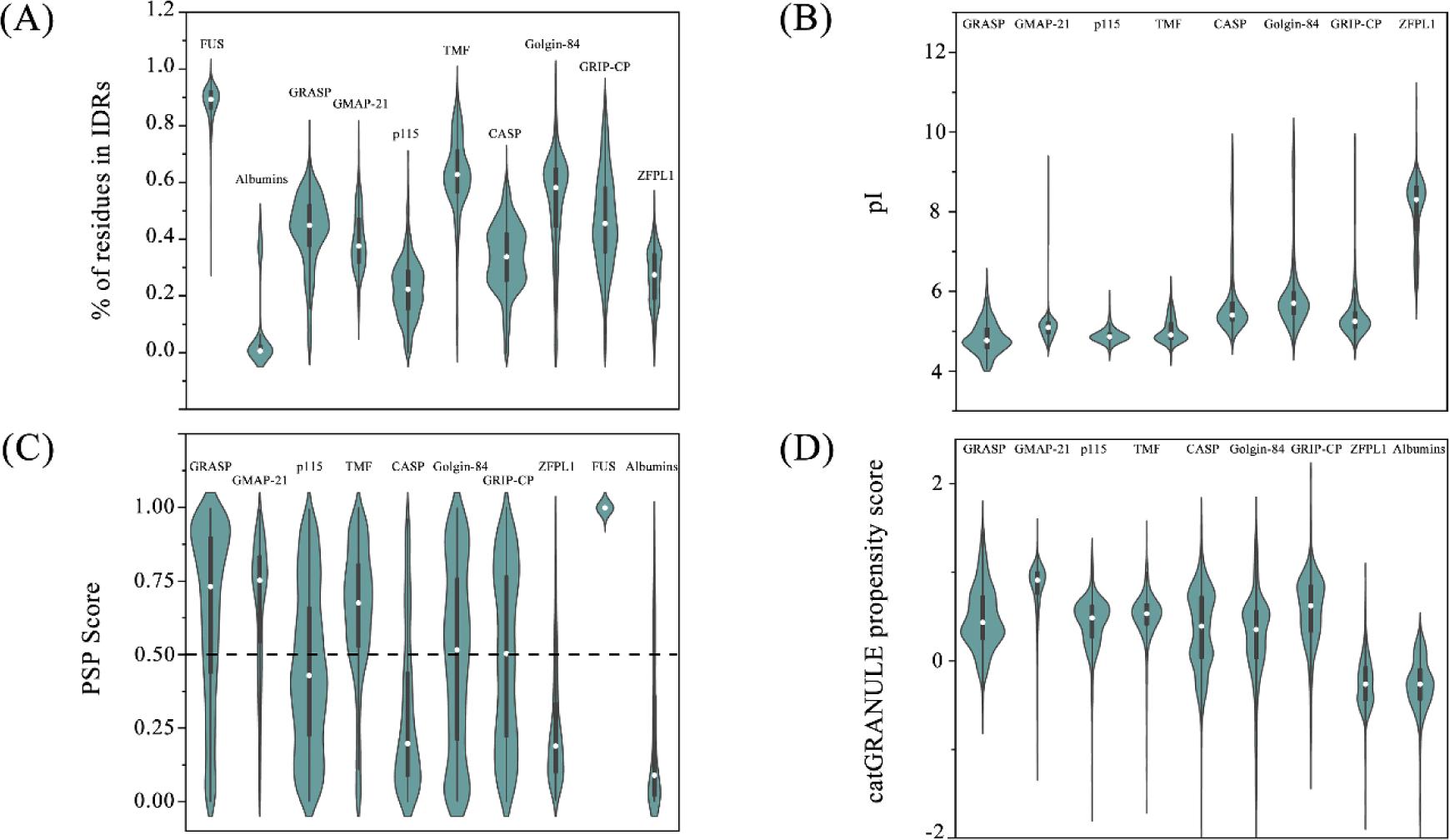
Analyses of the LLPS propensity and IDR content in Golgi structure proteins inferred to be present in the LECA according to (49). A) Representation of residue fractions exhibiting scores exceeding the 0.5 thresholds, using IUPred2 long disorder predictions. B) pI distribution across the GMP database. The pI values were calculated using an in-house Python script described in the Material and Methods Section. C) Application of the machine-learning-based PSPredictor (51). for LLPS prediction. A protein sequence is categorised as phase separating if the PsPred Score exceeds 0.5. D) Assessment of LLPS propensity utilising the catGRANULE algorithm, where propensity scores are evaluated in the context of proteome distribution, as conveyed through a cumulative distribution function. A higher score corresponds to a higher propensity for a protein to phase separate. As dosage-sensitive proteins are known to score approximately 0.5 (as per reference (52)), we designate this score as our threshold in the analyses.

Building on the detailed examination of GMPs’ theoretical propensity for LLPS, we now turn to experimental evidence to support our findings. To explore this phenomenon, we examined the physicochemical behaviour of human GRASPs in various solutions under crowded situations (58), aiming to understand and characterise the potential LLPS phenomena (Figure 3). Utilizing optical microscopy, we observed the formation of protein-rich droplets of GRASP55 (Figure 3A), and this tendency was both protein, ionic strength and crowding agent concentration-dependent (Figure S3). We initially focused on the impact of pH on the droplet formation. Considering the intracellular environment, we selected a pH of 5.5 to mimic extreme cellular conditions (59) and pH values of 6.0 and 6.3 to represent the values observed in *medial* and *trans* regions of the Golgi apparatus, respectively (60). Lastly, we included a physiological pH of 7.4 for comparative analysis. This approach allowed us to discern how varying pH conditions influence the behaviour of GRASP., contributing to our understanding of their role within the Golgi apparatus and potential implications for cellular function. Interestingly, despite GRASP65 being a vertebrate paralogue of GRASP55 (61), our experiments did not reveal any LLPS under the tested conditions (Figure 3B). The underlying factors contributing to this absence of LLPS in GRASP65 are not immediately apparent and warrant further investigation, which will be the subject of future investigation.

**Figure 3:**
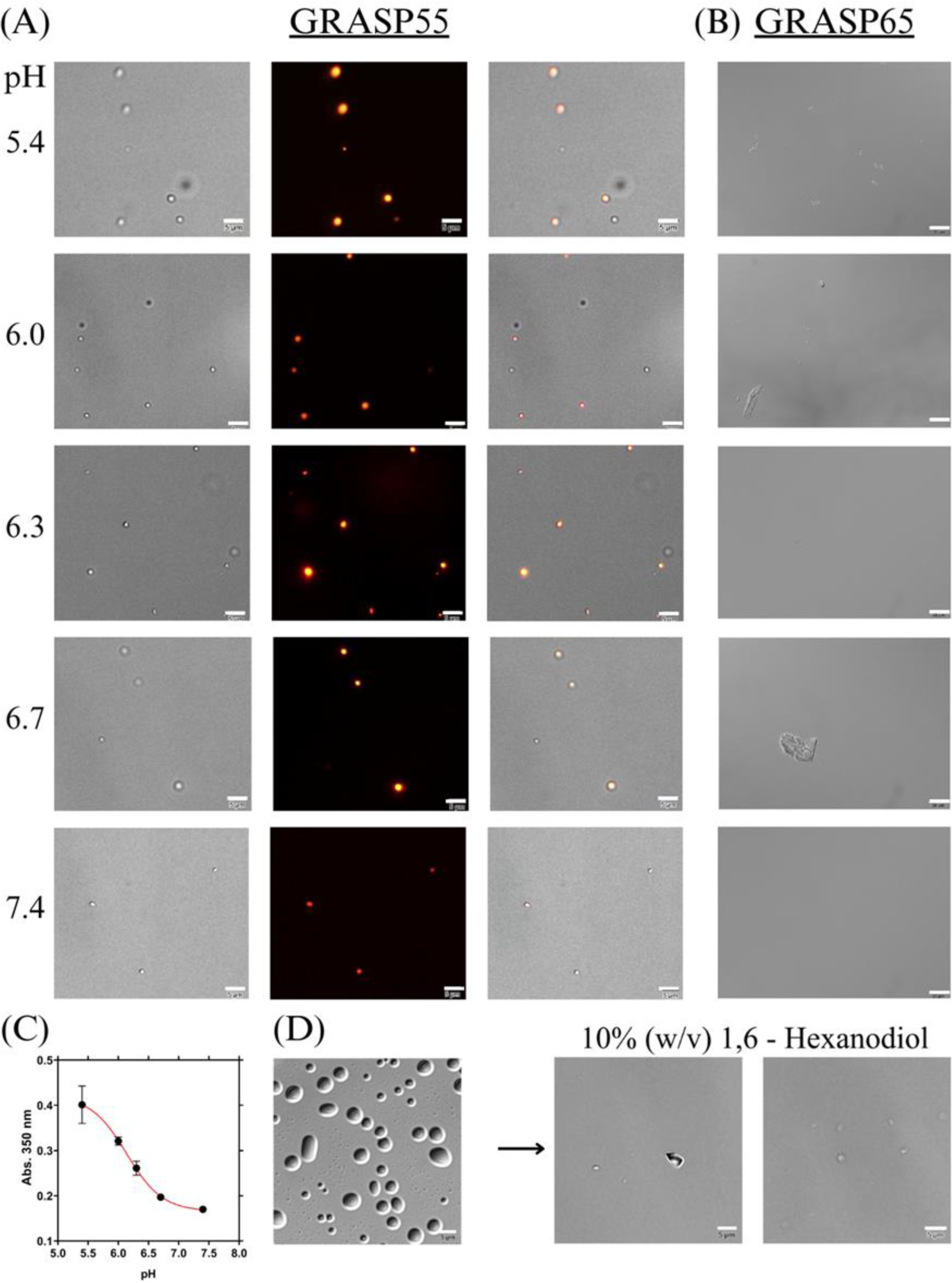
pH-Dependent Propensity of GRASPs for LLPS. (A) Differential Interference Contrast (DIC) microscopy images of GRASP55 (5 µM), highlighting the physical structure and morphology of the protein. The second row features fluorescence microscopy images of the same GRASP55 protein stained with Alexa Fluor 555 dye. (B) Similar to item (A) but using GRASP65. (C) Turbidity Analysis of GRASP55 phase separation at various pH levels, demonstrating pH-sensitive behaviour. (D) GRASP55 droplets dissolved after addition of 10% (w/V) 1,6 Hexanediol, monitored by DIC microscopy. The images selected are representative ones. Scale bars of 5 μm for (A) and (D), and of 20 μm for (B).

Our experiments revealed that GRASP55 underwent LLPS, although to a different extent, in all examined pH environments (Figure 3A). Turbidity assays showed that the most pronounced LLPS activity of GRASP55 occurred at pH levels corresponding to its native positioning within the *medial* (pH 6.0) and *trans* (pH 6.3) cisternae of the Golgi apparatus (Figure 3C) (62). This observation suggests a strong tendency for GRASP55 to engage in LLPS, potentially contributing to the structural and functional self-organisation within the Golgi complex.

We then established a pH of 6.3 for subsequent tests because it reflects the environment in the *medial* part of the Golgi apparatus, where GRASP55 demonstrated the most effective droplet formation. To further investigate the underlying forces driving LLPS in GRASP55, we conducted experiments using NaCl and 1,6-Hexanediol across a gradient range of 15 – 550 mM for NaCl and 0% - 15% for Hexanediol (Figure 3D and S3). The NaCl-containing samples demonstrated increased droplet formation with higher salt concentrations (Figure S3). In contrast, adding 10% Hexanediol led to the dissociation of the condensates after 10 minutes of incubation (Figure 3D). This result implies that the weak hydrophobic interactions disrupted by Hexanediol (63) are crucial in stabilising the GRASP55 droplets.

Besides examining the dependency of GRASP55 propensity to LLPS in a crowded environment, we also explored additional properties of the formed droplets, particularly their physical state – whether they were more liquid-like or gelatinous. We conducted permeability experiments using SYPRO™ Orange Protein Gel Stain to assess this. This stain permeates droplets if they are sufficiently liquid-like to allow for molecular exchange with their surrounding environment. In the initial part of our experiment, we prepared GRASP55 samples in LLPS-conditions and incubated them for 20 minutes to facilitate droplet formation. Subsequently, we checked the samples under a microscope to confirm the presence of droplets. Following this, we added SYPRO Orange to the solution at a concentration of 10% of the stock solution and incubated it for an additional 10 minutes to allow the SYPRO Orange to permeate the droplets. After 10 minutes of incubation, the samples were examined under a fluorescence microscope. Our observations revealed that the droplets exhibited liquid characteristics, as evidenced by the penetration of SYPRO Orange and subsequent fluorescence (Figure 4A). This liquid characteristic suggests that even after forming the biomolecular condensate, where the protein is at high concentrations, it can still exchange molecules with its surrounding environment. Another liquid-like observation in GRASP55 droplets was the occurrence of fusion or coalescence, as seen in Figure 4B, where individual protein droplets merged to form a single, larger droplet.

**Figure 4:**
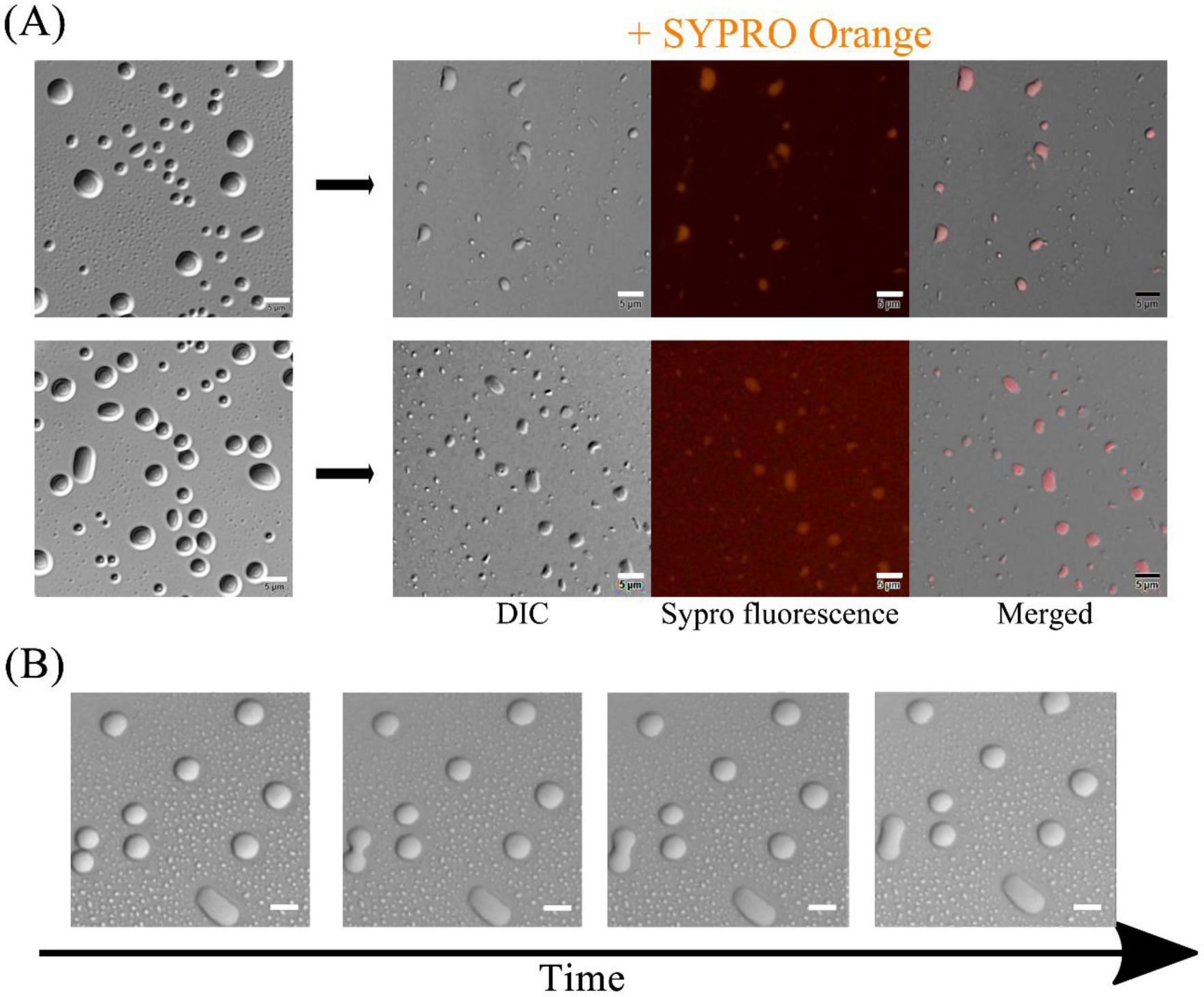
Liquid-like properties of GRASP55 droplets. (A) DIC Microscopy of GRASP55 (concentration of 5 µM). The image captures the dynamic process of droplets merging over time, illustrating the fluid nature of these structures. (B) The GRASP55 droplets (concentration: 5 µM) are observed under DIC microscopy. The image highlights the interaction of these droplets with the surrounding medium, where SYPRO Orange dye is dissolved. The permeation of the SYPRO Orange into the droplets is evident, indicating the liquid-like properties of the droplets and their ability to exchange substances with their environment—scale bars: 5 μm.

## Conclusion

In conclusion, our study provides a novel perspective on the nature of GMPs and critical functional characteristics across Eukarya, focusing on metazoans. Our data distinctly highlight the high prevalence of intrinsic disorder within GMPs, a feature that appears to be conserved across diverse eukaryotic lineages, albeit with different levels of prevalence and extent. Interestingly, the disorder in GMPs seems to be central to the potential of these proteins for LLPS, a phenomenon widely observed in the intracellular environment and critical for various biological processes. The combined findings indicate that disorder and LLPS propensity, despite the low primary sequence conservation across GMP families, have been evolutionarily conserved, suggesting their critical roles in the proteins’ functionality. To substantiate our findings, we conducted experimental tests on the potential for LLPS in a previously unstudied family of GMPs, namely GRASPs. These experiments confirmed their capacity for LLPS, thereby reinforcing our hypothesis. This study highlights that the functional versatility and adaptability of GMPs are likely derived from their structural disorder and/or elongated, multivalent properties. This intrinsic characteristic enables them to engage in diverse interactions and respond to various cellular signals, which is crucial for the organisation and maintenance of the Golgi apparatus. Our findings open the way for future studies on the role of LLPS in GMPs and their implication in various cellular processes. It will be particularly interesting to delve further into the precise mechanisms by which these proteins influence the architecture of the Golgi apparatus and the regulatory role of phase separation in these processes. The evolutionary conservation of these traits among GMPs suggests their fundamental importance in cell biology and invites further exploration.

## Material and Methods

### Database collection

In order to collect a broadly representative collection of unique protein orthologue sequences, we relied on the search for the protein family name in the hierarchical catalogue of orthologs OrthoDB (64) and for protein families in the InterPro database for those who are not catalogued in a single orthologue family (65). The protein sequences collected were as follows: TMF (OrthoDB 2054925at2759, 3717503at2759 and 1221595at2759, N = 1629), CASP (InterPro IPR012955, N = 2395), Golgin84 (InterPro IPR019177, N = 1500), Mammalian Serum Albumin-Like (from InterPro IPR020858, N = 1500), GRASP (OrthoDB 36330at2759, N = 2395), GM130 (OrthoDB 2880220at2759, N = 887), Golgin45 (OrthoDB 2879260at2759, N = 830), Golgin160 (OrthoDB 5357451at2759, N = 560), AKAP9 (OrthoDB 158920at33208, N = 1550), Giantin (OrthoDB 4227816at2759, N = 505), GCP60 (OrthoDB 38315at33208, N = 1384), GCC88 (OrthoDB 237371at33208, N = 831), Golgin97 (OrthoDB 42510at33208, N = 832), FUS RNA binding protein (OrthoDB 484351at7742, N = 982), Golgin245 (OrthoDB 2529099at33208, N = 871), protein bicaudal D homolog 2-like (OrthoDB 5350043at2759, N = 1886)), SCYL1BP1 (OrthoDB 2486431at33208, N = 708), p115 (OrthoDB 1774at2759, N = 1927), GCC185 (OrthoDB 814692at33208, N = 807), GRIP-Containing Proteins (N = 1500 randomly selected from InterPro IPR000237), GMAP (OrthoDB Group 5138860at2759, N = 886), ZFPL1 (OrthoDB Group 25738at2759, N = 903) and Janus kinase and microtubule interacting protein 2 - NECC (OrthoDB 5403492at2759, N = 1461). We employed the online resource found at https://fasta.bioch.virginia.edu/fasta_www2/clean_fasta.html (accessed in 2023) to process and clean our dataset’s FASTA formatted protein names. For most larger databases exceeding 1500 sequences, a custom Python script was utilised to randomly truncate the sequence data to the desired size of 1500 (script “fasta_random” in the SI). We have also generated a Python script to filter out unnatural amino acid residues within the protein sequences (script “fasta_filter” in the SI). This script reads the protein sequences from an input file, removes any character that is not a valid amino acid from each sequence, and writes the cleaned sequences to an output file, preserving the headers of the original sequences. This procedure was crucial given that most of the analysis tools we employed lacked the functionality to handle these unconventional letters appropriately. The full protein sequence database is available as a supplementary file.

### Prediction of LLPS-Propensity

The propensity for LLPS was evaluated using the PSPred and catGRANULE web-based platforms. Each algorithm accepts different sequence lengths: PSPred can simultaneously process segments of 100 amino acids, while catGRANULE can handle up to 500 amino acids per run. To streamline the upload process and accommodate these constraints, we developed Python scripts that partition our protein sequence database into appropriately sized segments for submission to each server. This script (script “fasta_split” in the SI) reads a FASTA file, splits the sequences into groups of a specified size, and writes each group to a separate FASTA file. The statistical analyses were performed using the software Origin 2022.

### Prediction of IDRs and calculated fractions within the sequence

We quantified each amino acid’s intrinsic disorder within a sequence using the IUPred2A tool, specifically designed for assessing long sequences (66). We engineered a Python script to interface with our comprehensive protein sequence databases to streamline the process (script “disorder_fractions” in the SI). This script seamlessly integrated with the IUPred2A software package we sourced directly from the official website. It then automated the computation of disorder scores, effectively isolating the fraction of residues with scores exceeding the threshold of 0.5. The statistical analyses were performed using the software Origin 2022.

### Charge vs Hydrophobicity Plot

We crafted a Python script implementing the Uversky methodology (script “fasta_CHA” in the SI) to systematically evaluate each protein sequence’s mean hydrophobicity and net charge in our database. Our user-friendly script allows customisation of the hydrophobicity scale per user preference, exploring the potential described in (67). For this manuscript, we have utilised the Kyte-Doolittle scale, originally employed by Uversky in (47). Our data presentation shows a discrimination borderline, defined by a specific equation, categorising proteins as either “natively disordered” or “natively ordered”. The equation is < *R* > = 2.785 < *H* > − 1.151, according to (47). In this work, < *H* > and < *R* > are the mean hydrophobicity and the mean net charge of the protein, respectively.

### Amino acid sequence general properties determination

We engineered a Python script to efficiently perform multiple tasks to expedite the protein sequence analysis process. This script reads protein sequences from a FASTA file, conducts a comprehensive evaluation of various properties using the Biopython package, and outputs these properties in an organised tabular format. The analysed properties include fractions of residues promoting order/disorder, isoelectric point (pI), Grand Average of Hydropathy (GRAVY), and the proportion of polar and charged residues. The statistical analyses were performed using the software Origin 2022.

### Protein expression and purification

GRASP55 and GRASP65 were purified as previously reported (68, 69). The purified solutions were concentrated with an Amicon Ultra-15 Centrifugal Filter (NMWL of 10 kDa, Merck Millipore, Burlington, USA). For protein concentration determination, the extinction coefficient at 280 nm was calculated with the ProtParam web server (70).

For the fluorescence microscopy experiments, protein samples were labelled with Alexa Fluor™ 555 NHS Ester (Succinimidyl Ester) (ThermoFisher). The dye was diluted in DMSO (Dimethyl sulfoxide pure) stock solution at a concentration of 10 mM. Before adding the dye, the protein solutions were exchanged with a 20 mM sodium phosphate buffer (Na2HPO4/NaH2PO4) at pH 7.4 using a HiTrap® Desalting Column connected to an Äkta purifier system (GE Healthcare). Subsequently, Alexa 555 dye was added at a concentration double that of the protein and the mixture was then incubated for approximately one hour at 4°C with gentle shaking. Post-incubation, excess unbound Alexa 555 was removed using a HiTrap® Desalting Column connected to an Äkta purifier system (GE Healthcare). Finally, the proportion of labelled to unlabelled protein was maintained at a 1% ratio for microscopy experiments.

### Differential interference contrast (DIC) and fluorescence microscopies

The LLPS and droplet formation by the target proteins were visualised using an inverted system microscope IX71 equipped with a PicoQuant MT 200 module (PicoQuant®, Germany) under DIC and fluorescence mode with a 60x objective (NA = 1.2). The fluorescence excitation intervals were selected using the mirror unities: U-MNU2 BP360/370; Blue: U-MWB2 BP460/490; Green: U-MWIG3 BP530-550. The long-pass filters HQ405LP, HQ460LP, BLP01-488R, HQ550LP and HQ690/70m were used. The droplet formations were induced by a 10X dilution method using a stock solution of 10 mg/mL unless otherwise stated. LLPS solutions were composed of 20 mM Na2HPO4/NaH2PO4 + X% (W/V) polyethylene glycol (PEG) 8000 (ThermoFisher) adjusted to the desired pH and with X varied from 2-20%. All the samples were incubated for 20 minutes in RT prior to measurements. The image acquisition was performed by adding 5uL drop solution in the centre of coverslips 20 x 20 mm (Deckgläser). Liquid droplet permeability was studied using 1X SYPRO™ Orange Protein Gel Stain (5,000X Concentrate in DMSO - Thermo Fisher Scientific). Data collection and analysis were performed with the SymPhoTime 64 software (PicoQuant).

### Turbidity assays

The turbidity experiments were conducted by measuring the absorbance at a wavelength of 350 nm in a Spectrophotometer Multiskan Go (ThermoFisher) equipment using the SkanIT RE 6.1.1 program. The samples contained only protein without any dye and the corresponding pH buffer in Na2HPO4/NaH2PO4 + X% (W/V) polyethylene glycol (PEG) 8000. The experiments were performed in triplicate with fresh sample preparation.

## Supporting information

Final protein sequences used

## Acknowledgements

The authors thank the *Fundação de Amparo à Pesquisa do Estado de São Paulo* (Grants No. 2022/06006-0, 2021/10465-7 and 2020/15542-7) for the financial support. AJCF thanks Conselho Nacional de Desenvolvimento Científico e Tecnlógico (CNPq) for the partial financial support (Grant no. 306682/2018-4)

## Supporting Information

In the supplementary material, we have included Figure S1-S3. Additionally, we have provided the Python scripts employed for data analysis. Furthermore, an accompanying .zip file contains the complete database referenced in our manuscript.

