## Supplementary Info for "Exploring Liquid-Liquid Phase Separation in the organization of Golgi Matrix Proteins"

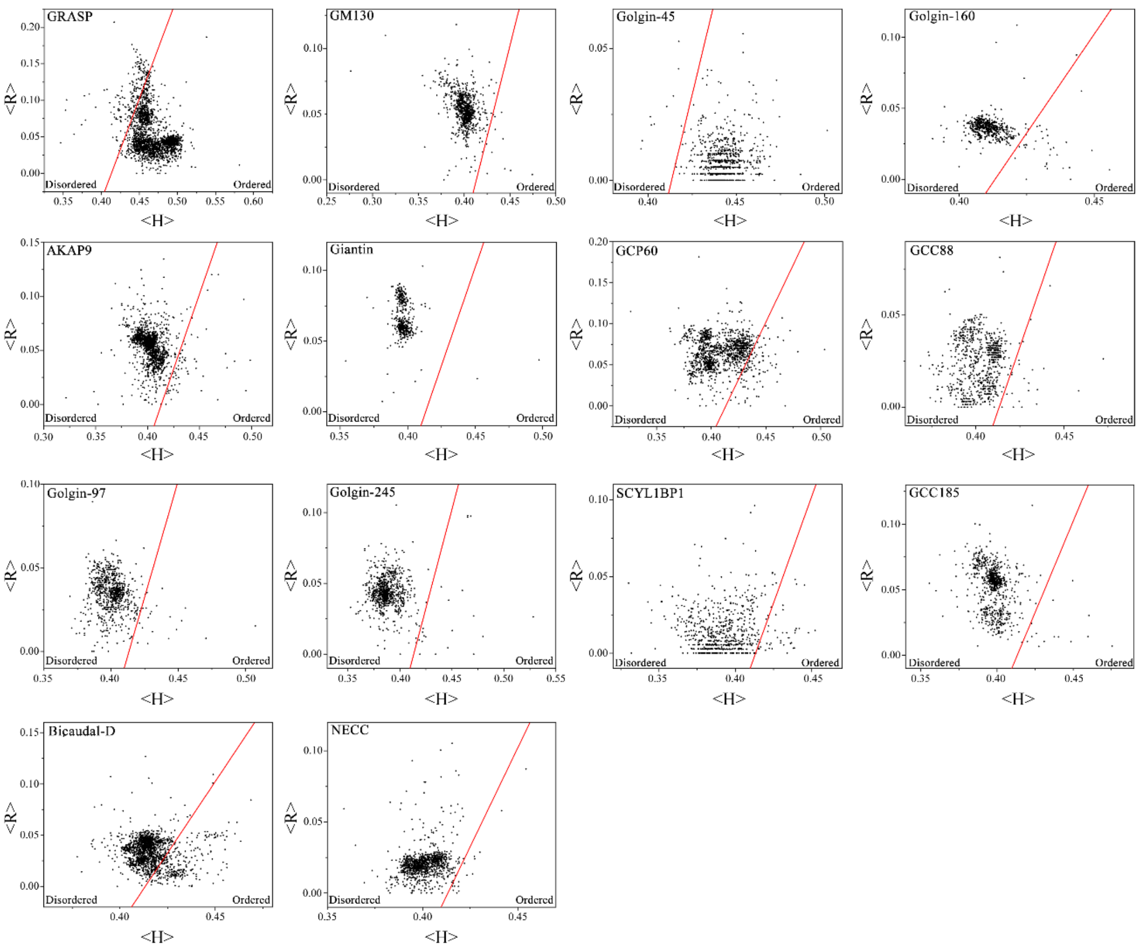


Figure S1: Charge-hydrophobicity phase space of the individual GMPs database where each black dot represents a specific GMP. The solid red line represents the border between "natively disordered" and "native ordered" proteins calculated according to the equation $< R > = 2.785 < H > \; - 1.151$. In this work, $< H >$ and $< R >$ are the mean hydrophobicity and the mean net charge of the protein, respectively. Both parameters and the borderline between ordered and disordered were calculated according to the Uversky methodology *(50)*. The number N of protein sequences ranges from ~500 to >2000 depending on the protein family, as presented in the material and methods section.

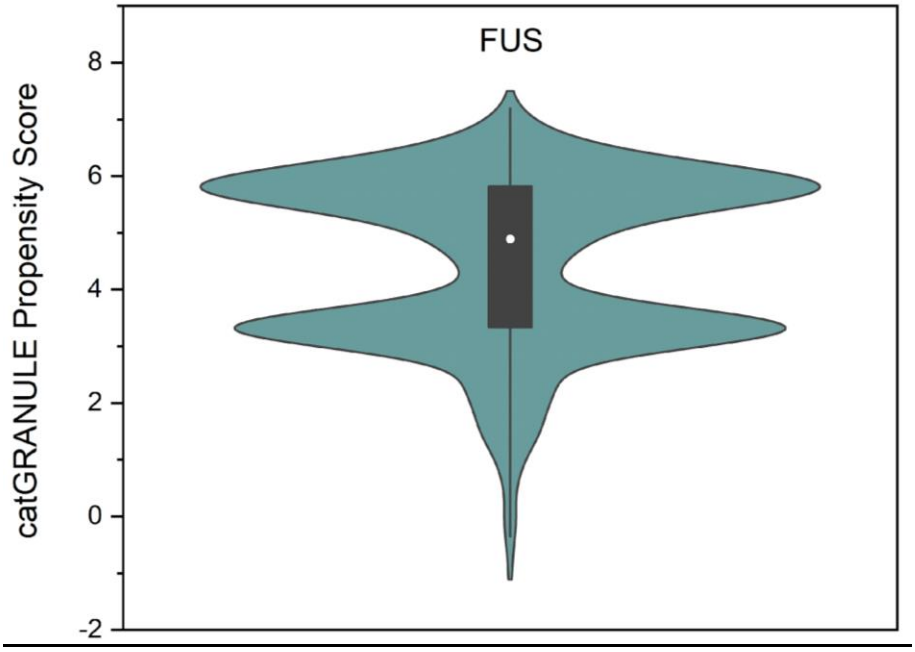


Figure S2: Assessment of FUS LLPS propensity utilising the catGRANULE algorithm, where propensity scores are evaluated in the context of proteome distribution, as conveyed through a cumulative distribution function. A higher score corresponds to a higher propensity for a protein to phase separate.

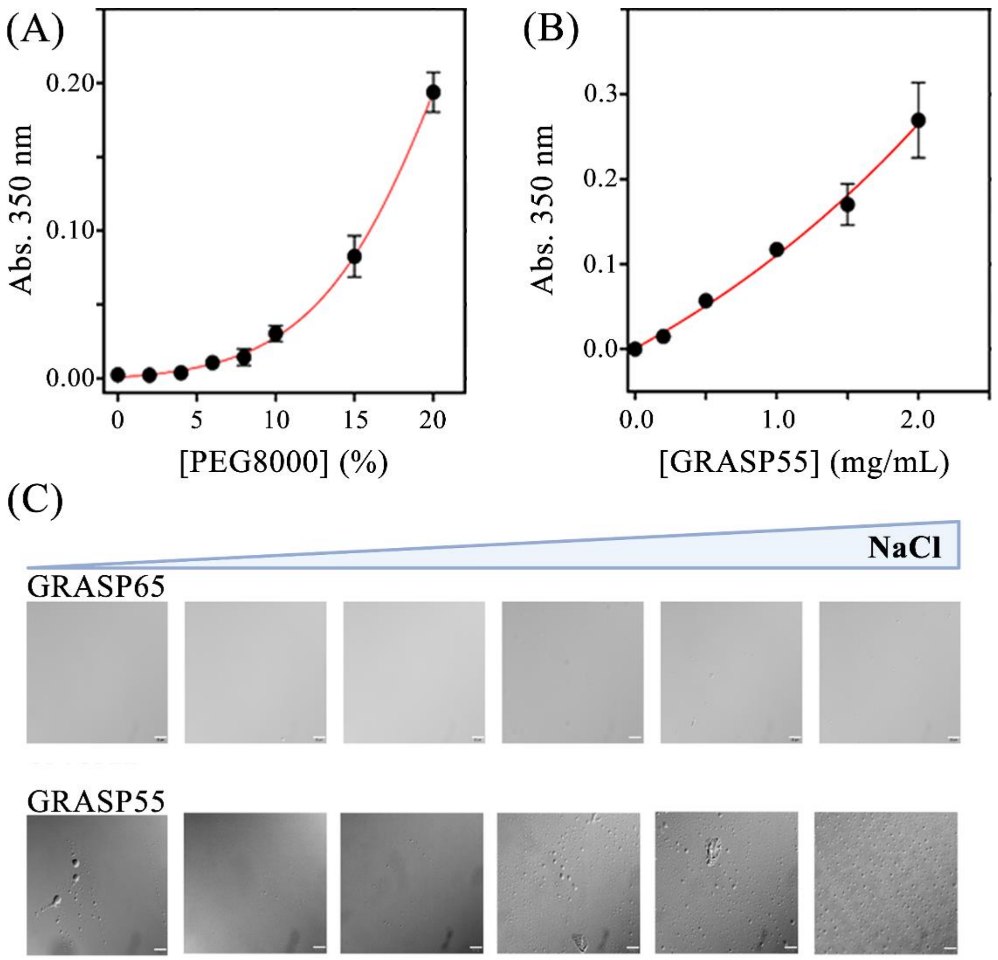


Figure S3: Liquid-liquid phase separation (LLPS) propensity of GRASP55/GRASP65. (A) Turbidity analysis showing GRASP55's phase separation behaviour across a PEG-8000 concentration gradient, illustrating PEG-dependent phase behaviour. (B) Turbidity analysis depicts the effect of GRASP55 concentration variations on phase separation to understand concentration impact further. (C) Differential Interference Contrast (DIC) microscopy images comparing LLPS phenomena: The top row features GRASP65 (5 µM), which does not undergo LLPS under any tested condition in the presence of NaCl, highlighting its distinct behaviour from GRASP55. The bottom row presents GRASP55 (5 µM) under NaCl conditions, demonstrating significant LLPS activity, thereby emphasizing the crucial role of electrostatic interactions in LLPS formation.

### List of Python Scripts for data analyses

*The original protein sequence database must be in fasta format. We have defined the input sequences to be listed in an input.txt file for all the scripts, using the Python 3.11 version.

### Script 1 – "fasta_filter"

```
# Define the list of valid amino acids
amino_acids = "ACDEFGHIKLMNPQRSTVWY"

# Open the input file in read mode
with open("input.txt", "r") as input_file:
    # Read all the records from the input file
    fasta_records = input_file.read().split(">")[1:]

    # Filter the records and remove non-amino acid characters from the sequences
    filtered_records = []
    for record in fasta_records:
        # Split each record into header and sequence
        header, sequence = record.strip().split("\n", 1)

        # Remove non-amino acid characters from the sequence
        sequence = "".join([aa for aa in sequence if aa in amino_acids])

        # Add the modified record to the list of filtered records
        filtered_records.append((header, sequence))

# Save the output file as "output.txt"
with open("output.txt", "w") as output_file:
    # Write the modified records to the output file
```

```
    for header, sequence in filtered_records:
        output_file.write(">" + header + "\n")
        output_file.write(sequence + "\n")
```

### Script 2 – "fasta_split"

```
def read_fasta(filename):
    """Reads a FASTA file and returns a list of tuples with the header and sequence."""
    sequences = []
    with open(filename, "r") as f:
        header = ""
        seq = ""
        for line in f:
            if line.startswith(">"):
                if seq:
                    sequences.append((header, seq))
                    seq = ""
                header = line.strip()
            else:
                seq += line.strip()
        sequences.append((header, seq))
    return sequences

def write_fasta(sequences, filename):
    """Writes a list of sequences to a FASTA file."""
    with open(filename, "w") as f:
        for seq in sequences:
            f.write(seq[0] + "\n" + seq[1] + "\n")
```

```
def split_sequences(sequences, max_per_file):
    """Splits a list of sequences into several lists with a maximum of max_per_file sequences."""
    split_sequences = []
    current_seqs = []
    for seq in sequences:
        current_seqs.append(seq)
        if len(current_seqs) == max_per_file:
            split_sequences.append(current_seqs)
            current_seqs = []
    if current_seqs:
        split_sequences.append(current_seqs)
    return split_sequences

if __name__ == "__main__":
    # Define the name of the input file and the maximum number of sequences per output file
    input_file = "input.txt"
    max_per_file = 500

    # Read the sequences from the input file
    sequences = read_fasta(input_file)

    # Splits the sequences into several groups, each with at most max_per_file sequences
    split_sequences = split_sequences(sequences, max_per_file)

    # Writes each group of sequences to a separate FASTA file
    for i, seq_group in enumerate(split_sequences):
        output_file = "sequences_{}.fasta".format(i+1)
        write_fasta(seq_group, output_file)
```

### Script 3 – “fasta_random”

```
import random


# Input file name
input_file = "input.txt"
# Output file name
output_file = "output.txt"
# Number of sequences to be selected
num_sequences = 1500


# Opens the input file and reads all sequences into a list
with open(input_file, "r") as f:
    sequences = f.read().split(">")[1:]


# Randomly selects sequences
selected_sequences = random.sample(sequences, num_sequences)


# Writes the selected sequences to the output file
with open(output_file, "w") as f:
    for seq in selected_sequences:
        f.write(f">{seq}")
```

### Script 4 – “fasta_CHA”

```
import re
from Bio import SeqIO
from Bio.SeqUtils.ProtParam import ProteinAnalysis


# Huang et al. BMC Bioinformatics 2014, 15(Suppl 17):S4
GUY_SCALE = {
```

```
    "A": 0.1, "R": 1.91, "N": 0.48, "D": 0.78, "C": -1.42,
    "Q": 0.83, "E": 0.95, "G": 0.33, "H": -0.5, "I": -1.18,
    "L": -1.8, "K": 1.4, "M": -1.59, "F": -2.12, "P": 0.73,
    "S": 0.52, "T": 0.07, "W": -0.51, "Y": -0.21, "V": -1.27
}

KYTE_DOOLITTLE_SCALE = {
    "A": 1.8, "R": -4.5, "N": -3.5, "D": -3.5, "C": 2.5,
    "Q": -3.5, "E": -3.5, "G": -0.4, "H": -3.2, "I": 4.5,
    "L": 3.8, "K": -3.9, "M": 1.9, "F": 2.8, "P": -1.6,
    "S": -0.8, "T": -0.7, "W": -0.9, "Y": -1.3, "V": 4.2
}

IDP_HYDROPATHY_SCALE = {
    "A": 0.91, "R": 0.07, "N": 2.06, "D": -0.48, "C": 5.62,
    "Q": -1.23, "E": -2.20, "G": 0.02, "H": 2.18, "I": 6.19,
    "L": 5.17, "K": -2.43, "M": 2.49, "F": 5.79, "P": -3.89,
    "S": -1.84, "T": 1.22, "W": 10.66, "Y": 6.64, "V": 4.64
}

def normalize_hydrophobicity(seq, scale):
    normalized = []
    for aa in seq:
        if scale == "Guy":
            value = GUY_SCALE[aa]
            min_value = min(GUY_SCALE.values())
            max_value = max(GUY_SCALE.values())
            # Invertendo a normalização da escala Guy
            normalized_value = 1 - ((value - min_value) / (max_value - min_value))
        elif scale == "Kyte-Doolittle":
            value = KYTE_DOOLITTLE_SCALE[aa]
```

```
            min_value = min(KYTE_DOOLITTLE_SCALE.values())
            max_value = max(KYTE_DOOLITTLE_SCALE.values())
            normalized_value = (value - min_value) / (max_value - min_value)
        else:
            value = IDP_HYDROPATHY_SCALE[aa]
            min_value = min(IDP_HYDROPATHY_SCALE.values())
            max_value = max(IDP_HYDROPATHY_SCALE.values())
            normalized_value = (value - min_value) / (max_value - min_value)
        normalized.append(normalized_value)
    return normalized


def analyze_sequence(filename, scale):
    with open(filename) as file:
        records = SeqIO.parse(file, "fasta")
        result = []
        for record in records:
            seq = str(record.seq)
            analysis = ProteinAnalysis(seq)
            num_residues = len(seq)
            total_neg = analysis.count_amino_acids()['D'] + analysis.count_amino_acids()['E']
            total_pos = analysis.count_amino_acids()['R'] + analysis.count_amino_acids()['K']
            if scale == 1:
                hydrophobicity_normalized = normalize_hydrophobicity(seq, "Guy")
            elif scale == 2:
                hydrophobicity_normalized = normalize_hydrophobicity(seq, "Kyte-Doolittle")
            else:
                hydrophobicity_normalized = normalize_hydrophobicity(seq, "IDP-Hydropathy")
            hydrophobicity_avg = sum(hydrophobicity_normalized) / num_residues
            mean_net_charge = abs((total_pos - total_neg) / num_residues)
            result.append((record.id, mean_net_charge, hydrophobicity_avg))
        return result
```

```
def write_output(filename, results):
    with open(filename, "w") as file:
        file.write("ID\tMean_Net_Charge\tMean_Hydrophobicity\n")
        for r in results:
            file.write(f"{r[0]}\t{r[1]:.4f}\t{r[2]:.4f}\n")


if __name__ == "__main__":
    input_file = "input.txt"
    output_file = "output.txt"
    scale = 0
    while scale not in [1, 2, 3]:
        scale = int(input("Choose the hydrophobicity scale: type '1' for Guy, '2' for Kyte-Doolittle or '3' for IDP-Hydropathy: "))
    results = analyze_sequence(input_file, scale)
    write_output(output_file, results)
```

### Script 5 – "protein_prop"

```
import sys
from Bio import SeqIO
from Bio.SeqUtils.ProtParam import ProteinAnalysis
from tabulate import tabulate
import pandas as pd


# Lists of amino acids types
hydrophobic = ['A', 'V', 'I', 'L', 'M', 'F', 'Y', 'W', "G"]
charged = ['D', 'E', 'R', 'H', 'K']
polar = ['Q', 'N', 'P', 'S', 'T', 'C']
```

```
def read_fasta_file(input_file):
    fasta_sequences = []
    with open(input_file, "r") as file:
        for record in SeqIO.parse(file, "fasta"):
            fasta_sequences.append(record)
    return fasta_sequences

def calculate_pI(protein_sequence):
    analysis = ProteinAnalysis(str(protein_sequence))
    pI = analysis.isoelectric_point()
    return pI

def calculate_molecular_weight(protein_sequence):
    analysis = ProteinAnalysis(str(protein_sequence))
    molecular_weight = analysis.molecular_weight()
    return molecular_weight

def calculate_GRAVY(protein_sequence):
    analysis = ProteinAnalysis(str(protein_sequence))
    GRAVY = analysis.gravy()
    return GRAVY

def calculate_fraction(seq, list):
    count = 0
    for aa in str(seq):
        if aa in list:
            count += 1
    return count / len(seq)

def write_output(output_file, protein_properties):
    with open(output_file, "w") as file:
```

```
        table = tabulate(protein_properties, headers=["Protein Name", "Molecular Weight", "pI", "GRAVY", "Fraction Hydrophobic", "Fraction Charged", "Fraction Polar"], tablefmt="pipe")
        file.write(table)


def main():
    input_file = "input.txt"
    output_file = "output_protein_properties.txt"

    fasta_sequences = read_fasta_file(input_file)
    protein_properties = []

    for sequence in fasta_sequences:
        pI = calculate_pI(sequence.seq)
        molecular_weight = calculate_molecular_weight(sequence.seq)
        GRAVY = calculate_GRAVY(sequence.seq)
        fraction_hydrophobic = calculate_fraction(sequence.seq, hydrophobic)
        fraction_charged = calculate_fraction(sequence.seq, charged)
        fraction_polar = calculate_fraction(sequence.seq, polar)

        protein_properties.append([sequence.id, molecular_weight, pI, GRAVY, fraction_hydrophobic, fraction_charged, fraction_polar])

    write_output(output_file, protein_properties)

if __name__ == "__main__":
    main()
```

### Script 6 – "disorder_fractions"

```
import os
import subprocess
```

```
from Bio import SeqIO
from statistics import mean

def iupred2a_local(sequence, iupred2a_script_path, iupred2a_type="long"):
    with open("temp_sequence.fasta", "w") as temp_file:
        temp_file.write(">temp_sequence\n" + sequence)

    command = f'python "{iupred2a_script_path}" temp_sequence.fasta {iupred2a_type}'
    result = subprocess.check_output(command, shell=True).decode("utf-8")
    os.remove("temp_sequence.fasta")

    disordered_scores = []
    for line in result.split("\n"):
        if not line.startswith("#"):
            columns = line.split()
            if len(columns) >= 3:
                try:
                    score = float(columns[2])
                    disordered_scores.append(score)
                except ValueError:
                    pass

    return disordered_scores


def calculate_disorder_fraction(sequence, iupred2a_script_path):
    disordered_scores = iupred2a_local(sequence, iupred2a_script_path)
    threshold = 0.5
    disordered_count = sum(1 for score in disordered_scores if score > threshold)
    return disordered_count / len(sequence)
```

```
def main(fasta_file, iupred2a_script_path):
    disorder_fractions = []

    with open(fasta_file, "r") as file:
        fasta_records = list(SeqIO.parse(file, "fasta"))
        for record in fasta_records:
            sequence = str(record.seq)
            disorder_fraction = calculate_disorder_fraction(sequence, iupred2a_script_path)
            disorder_fractions.append(disorder_fraction)

    avg_disorder_fraction = mean(disorder_fractions)

    with open("output.txt", "w") as output_file:
        for disorder_fraction in disorder_fractions:
            output_file.write(f"{disorder_fraction}\n")
        output_file.write(f"\nAverage fraction of disordered residues: {avg_disorder_fraction}\n")

    print("The results were saved in output.txt.")


if __name__ == "__main__":
    fasta_file = r"C://input.txt"
    iupred2a_script_path = r"C://iupred2a.py"
    main(fasta_file, iupred2a_script_path)
```
